# Homogeneous large field-of-view and compact iSCAT-TIRF setup for dynamic single molecule measurements

**DOI:** 10.1101/2024.06.18.599482

**Authors:** Giovanni De Angelis, Jacopo Abramo, Mariia Miasnikova, Marcel Taubert, Christian Eggeling, Francesco Reina

## Abstract

Interferometric Scattering Microscopy (iSCAT) enables prolonged and high frame rate Single Particle Tracking (SPT) for single molecule dynamics studies. Typically, iSCAT setups employ scanning illumination schemes to achieve uniform sample illumination. However, this implementation limits the field of view (FoV) and maximum sampling rate, while increasing hardware requirements and setup size. We demonstrate the realization of a large (60µm x 60µm) uniformly illuminated FoV through a passive refractive optical element in the iSCAT illumination path. This scanning-free iSCAT microscopy setup is further combined with an objective based Total Internal Reflection Fluorescence Microscopy (TIRF) channel for a complementary fluorescence readout, a focus-lock system, and a tailored control platform via the open-source ImSwitch software, and has a compact footprint. As a proof-of-principle, we highlight the performance of the setup through the acquisition of iSCAT images with a uniform contrast and a ≤10 nm localization precision throughout the whole FoV. The performance is further demonstrated through dynamic iSCAT SPT and imaging Fluorescence Correlation Spectroscopy of lipid diffusion in a model membrane system. Our iSCAT setup thus depicts an accurate and improved way of recording fast molecular dynamics in life sciences.

## 1. Introduction

Interferometric Scattering (iSCAT) microscopy is a relatively novel technique that has shown great potential since its first developments for single-molecule measurements [1–7]. The main favorable characteristic of iSCAT microscopy stems from its image formation process. In iSCAT, the contrast is derived from the interference between a reference beam, originated by the reflection of the incoming coherent illumination at the interface between the sample media and its glass support, and the component of the light incident on the sample backscattered by any object whose refractive index is different from that of the surrounding medium [8]:

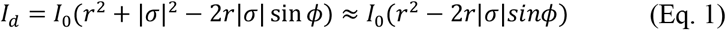

where *I*_*d*_ is the detected intensity at the sensor, *I*_0_ is the light incident at the sample, *r* is the reflectivity of the interface between the sample medium and the glass support, *σ* is the scattering cross section of an object in the sample, and *ϕ* is the phase difference between the two beams. The last part of (Eq. 1) omits the pure scattering term, as the scattering cross section is typically much smaller than the reflectivity *r*. The linear dependence of the detected signal from the scattering cross section is the main reason behind the increased sensitivity of iSCAT compared to scattering based technique such as dark field microscopy [1]. This translates directly to the exceptional sensitivity of iSCAT to the scattering of nanoscopic objects, such as single proteins, which has been recently exploited successfully to perform so-called mass photometry [9–11]. This high sensitivity, together with the reliance of scattering for image formation that affords higher sampling rates than fluorescence-based methods, have further solidified the significance of iSCAT microscopy as a single molecule-sensitive technique with exceptionally good performance for the observation of molecular diffusion dynamics in cells and membranes via single particle tracking (SPT) [12–17].

From a purely technical standpoint, iSCAT microscopy is of relatively simple implementation in a laboratory, provided good expertise with optical techniques is available. One of the most common implementations of iSCAT involves using the coherent illumination light in widefield mode, which emphasizes the imaging sampling rate of the device. An important feature thereby is to realize a rather uniform illumination profile to entail homogeneous scattering signal and thus statistics over the whole field-of-view (FoV). The most straightforward approach is a conventional wide-field illumination with a Gaussian beam and thus Gaussian focal intensity profile. However, this leads to accurate uniformity only within a small region around the peak of the Gaussian intensity profile, generating a rather small FoV. Further, and more important to iSCAT microscopy, is the generation of time-variable interference fringes and speckle noise in the iSCAT signal that deteriorate image quality, especially when conjugated with high-NA objectives [8,18].

A well laid-out protocol by Ortega-Arroyo and coworkers has provided the community with useful instructions and indications for the construction of an optical setup that is capable of iSCAT microscopy with both uniform and interferences- and speckle-free illumination [18]. The setup described in that protocol features an illumination line with two acousto-optic deflector scanners (AODs) for raster scanning the sample. When the scattered light is collected by the camera sensor from several scanning cycles, and the AOD scanning and the camera detection are synchronized, this allows the sample to appear uniformly illuminated. This approach has therefore been implemented in multiple applications. Nevertheless, this detection scheme presents some disadvantages that could hinder the widespread adoption of iSCAT. For example, the large footprint implied in the realization of this setup poses a significant challenge for environments with space constraints. Moreover, the alignment and synchronization of two AOD scanners can be somewhat challenging. Finally, scanning of the sample naturally enhances data acquisition time and thus may reduce time resolution of an iSCAT experiment.

For these reasons, we explored the possibility of guaranteeing uniform illumination across a large field-of-view in iSCAT detection without beam scanning. This essentially means implementing widefield illumination, as it would be necessary to focus the incident light on the back focal plane of the microscope objective. As highlighted before, several literature sources indicate disadvantages of such a procedure, for example the generation of time-variable interference fringes and speckle noise that deteriorate image quality, especially when conjugated with high-NA objectives [8,18–24]. However, if the setup is interferometrically stable, such contributions should be constant in time and thus may be subtracted as background, like it is commonly the case in iSCAT microscopy [3,6,18,20]. Given that a typical iSCAT setup is, essentially, a common-path interferometer, we saw no *a priori* difficulty in achieving the required stability to implement a widefield illumination scheme. Further, many designs for a homogeneous intensity distribution across a large FoV have been realized throughout the years, in particular to improve the performance of single molecule localization fluorescence microscopy, i.e. to guarantee a comparable localization precision across the FoV [25–27]. Our choice here is a commercial device which shapes a collimated Gaussian beam to a collimated beam of uniform irradiance, that is, a top-hat intensity profile [28]. This is accomplished by using a system of plano-aspheric lenses, which ensures that the desired properties of the illumination, i.e. collimation and uniform irradiance, are preserved over a sufficiently long distance [29–31]. Such a device has already been applied successfully to single-molecule fluorescence microscopy [27], but not yet to scattering-based microscopy techniques. Equivalent illumination schemes should however lead to similar performance [32].

In this work, we show that our microscope design, with a flat intensity widefield illumination scheme, performs adequately in all the applications for which iSCAT microscopy is desirable, i.e. imaging, SPT and complementary fluorescence recordings. The optical setup features simpler implementation and reduced footprint compared to similar devices reported in literature. We further demonstrate its compatibility with Total Internal Reflection Fluorescence (TIRF) microscopy for correlative fluorescence recordings, and how focus stabilization can be implemented by optical means. Finally, we provide guidance on how to control the microscope through an open-source software (ImSwitch), which may be rendered compatible with the majority of commercially available components.

## 2. Results and Discussions

### 2.1 iSCAT setup

We realized an iSCAT microscope design with a flat intensity widefield illumination scheme, a small instrument footprint (<1m^2^), the possibility for correlative TIRF microscopy recordings and focus stabilization. The resulting scanning-free and large FoV was capable of imaging large spanning sample regions (∼60×60 µm^2^) without tiling. The basic setup is depicted in Fig. 1 and described in detail in the Methods section.

**Fig. 1.**
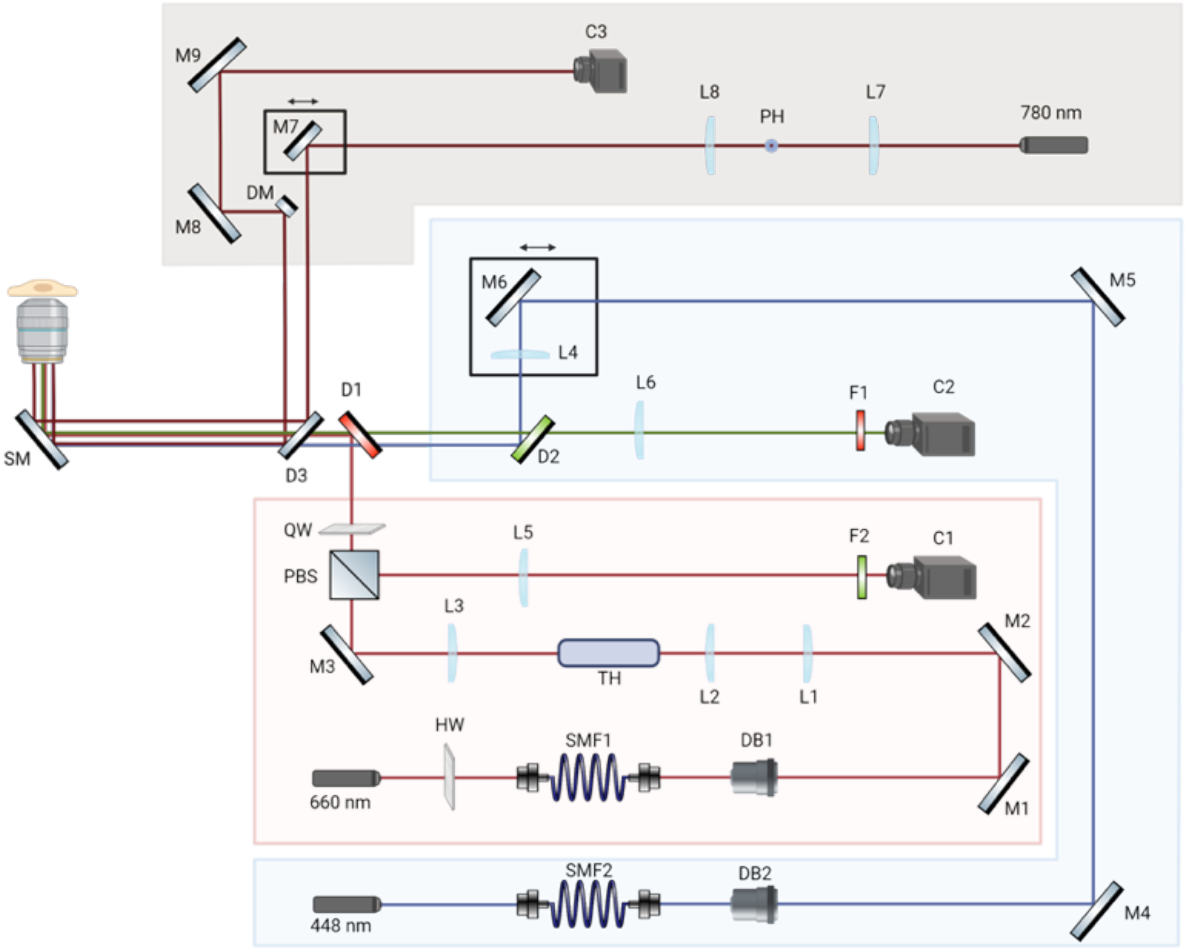
iSCAT-TIRF setup. Red box = iSCAT channel. Blue box = TIRF Channel. Brown box = focus-lock channel. SMF1-SMF2, single mode fiber. HW, half-wave plate. QW, quarter-wave plate, M1-M9, dielectric mirror. SM, silver mirror. PH, pin holes. DB1-DB2, dry objectives. L1-L8 plano-convex lenses. D1-D3, dichroic mirrors. F1-F2, single notch filters. C2, sCMOS cameras. C1-C3, CMOS camera.

While implementing a conventional widefield illumination through focusing into the back focal plane of the microscope objective lens, scattering-based techniques encounter inhomogeneous features in the raw images. This is due to several factors such as nonuniform illumination, spurious back reflections and unwanted interferences caused by reflections or defects of optical elements [18]. Filtering out these intrinsic features in post-processing is only possible if the setup is interferometrically stable, meaning that the interference or speckle patterns do not change in the experimental time frame, as is highlighted for our setup in Fig. S1. To avoid the intrinsic illumination inhomogeneity caused by a Gaussian beam illumination, we specifically used a refractive optical component to shape the beam into a Top Hat (TH) beam profile [27]. We then focused the TH beam onto the back focal plane of a high NA objective (Materials and Methods) leading to a homogeneous sample illumination. The scattering signal was then detected with a fast CMOS camera with 1280 × 864 pixels and up to > 3.5 kHz frame rate. Fig. S1 highlights the interferometric stability of our setup and that the interference patterns were turned into a homogeneous TH intensity profile after processing of the data, which is detailed and highlighted further below.

In addition to the iSCAT channel, we implemented a fluorescence channel, able to switch between epi- and total internal reflection (TIRF) illumination, similar to what was shown in [18]. This fluorescence channel enabled correlative imaging and single-molecule sensitive detection like imaging fluorescence correlation spectroscopy (imaging FCS) [33–36], as described later on.

Also, we added a focus-lock system similar to [37], which monitors the displacement of a totally internal reflected 780 nm laser from the cover glass. Repeated imaging of gold beads immobilized the cover glass highlighted a focal drift along the axial z-direction of less than 65 nm over 6 seconds.

The instrument control, data acquisition as well as basic data processing were custom-written in ImSwitch, a generic software for microscope instrumentation [38] (see Materials and Methods).

### 2.2 iSCAT image processing

The image formation of the scattered signal in iSCAT leads to strong background contributions in the camera image (Fig. 2). This was due to multiple reflections generated by the several optical components within the optical system, which created a static interference pattern in the image that was not related to the sample (Fig. S1). This is a well-known phenomenon in iSCAT microscopy, which can straightforwardly be corrected with a temporal median filter [1,39]. Thus, we implemented a real-time median filter correction that was applied on the fly to the raw image directly in ImSwitch [38]. This was specifically realized through a synchronization mechanism which we provided as part of the control software. This enabled straightforward evaluation of the sample in real time. The median filtering procedure as integrated into our software was thus extremely simple. The sample was automatically moved by the motorized stage by a fixed number of steps (see Methods), and an image was acquired at each step. The software then generated a temporal median filter from the acquired image stack. The live images were then divided by the median filter leading to a background-free image where the scattering objects (40nm gold nanoparticles) could accurately be identified as dark spots (Fig. 2).

**Fig. 2.**
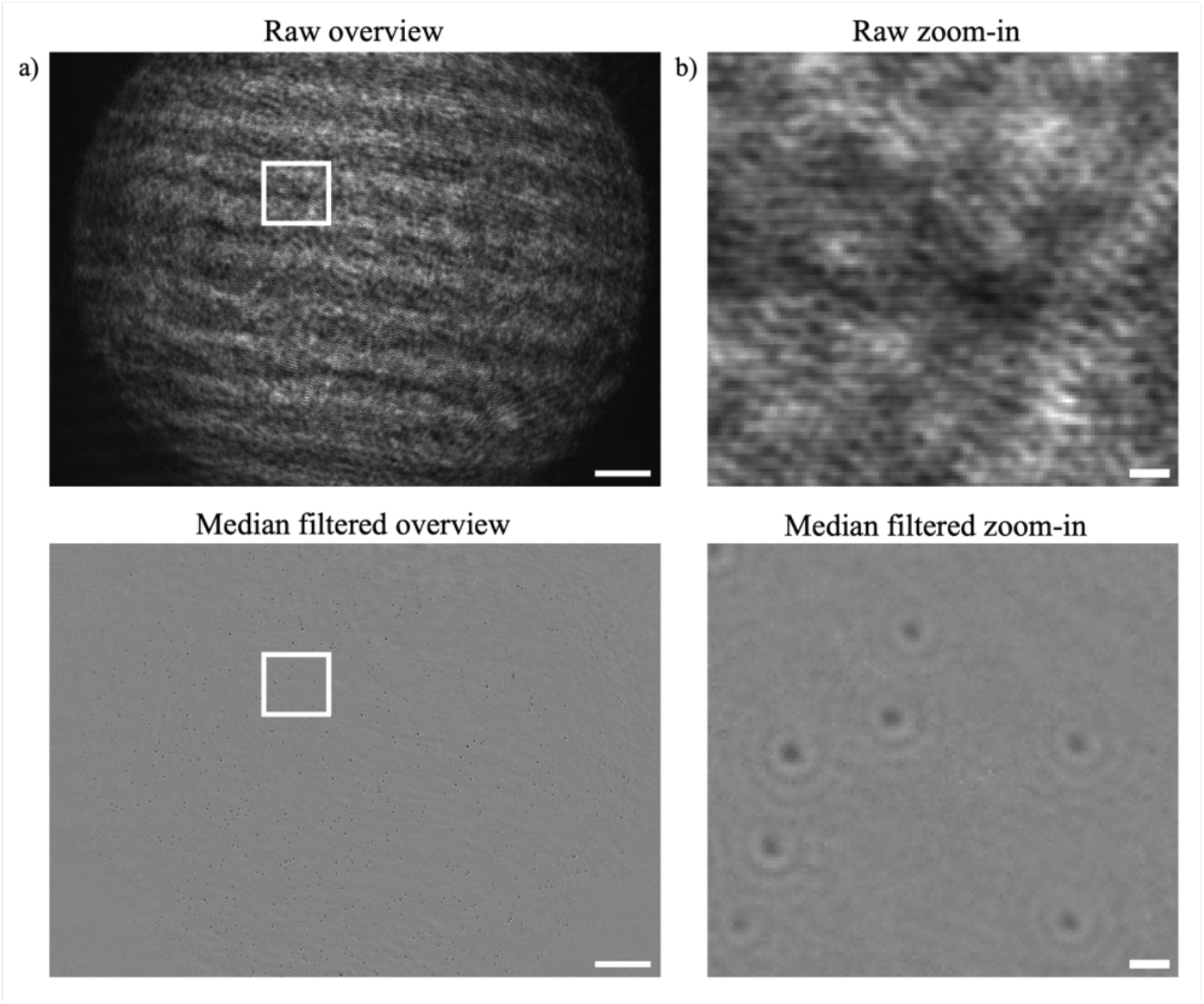
iSCAT image data processing. (a) Representative full chip field of views of the iSCAT images before (upper panel) and after (lower panel) the application of the median filter. Scale bar 10 *µm*. (b) Close-up images of the same FoV region (white boxed in (a)) before (upper panel) and after (lower panel) the application of the median filter, revealing the presence of 40nm gold nanoparticles attached to the coverglass as dark spots with interference fringes. Scale bar 1 *µm*.

### 2.3 Characterization of the uniform illumination profile

Homogeneous sample illumination has multiple advantages with respect to a Gaussian illumination intensity profile. As shown in Fig. 3a, the FoV achieved with our homogeneous TH illumination arrangement was 60×60 *µm*^2^, which was wider and more homogeneous than what was obtained with a Gaussian illumination with a full width at half maximum (FWHM) of 28 *µm*. To measure the latter, we removed the refractive TH component and focused the laser beam directly into the back focal plane of the microscope objective. Here it is important to highlight that the only difference between the two illumination arrangements was indeed the presence of the TH component, and no other optical components were changed in the setup to perform the comparison.

**Fig. 3.**
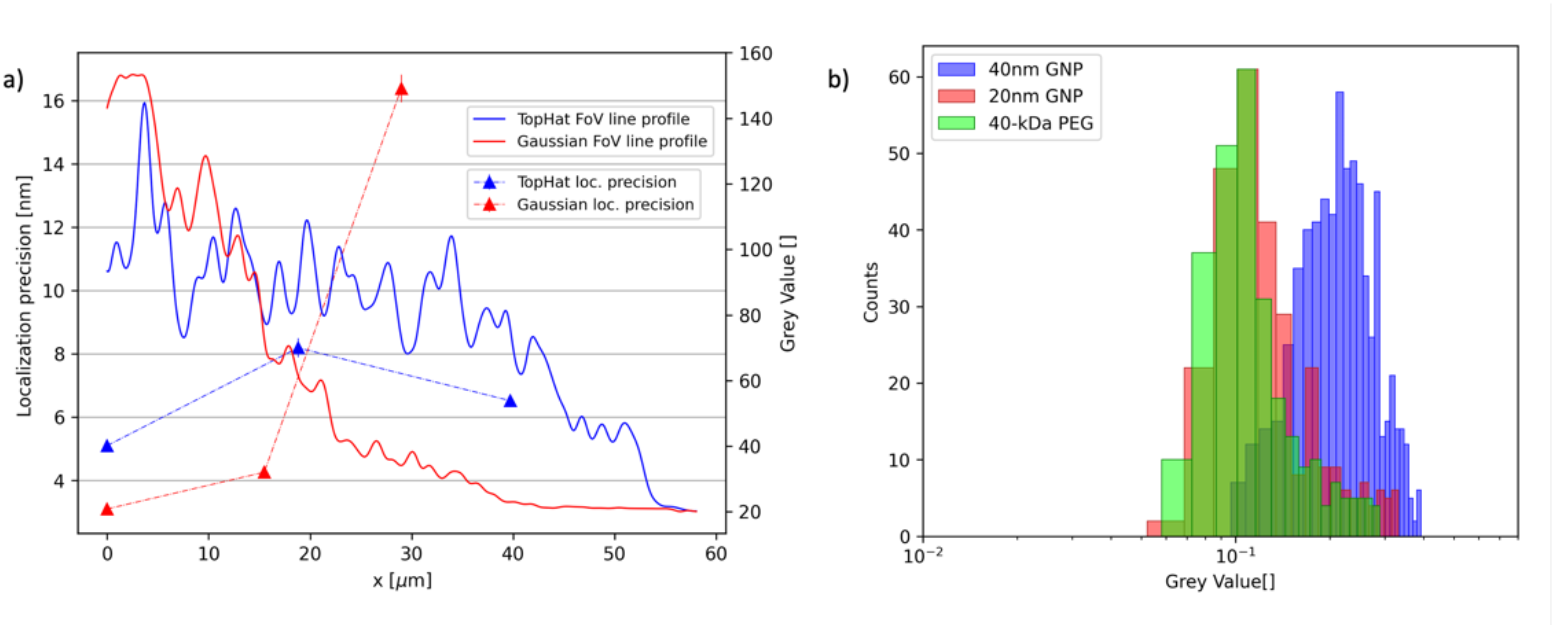
Homogeneity and accuracy of the iSCAT recordings over the whole field-of-view (FoV). (a) Comparison between the homogeneous top hat and Gaussian illumination profile: laser intensity line profile through the FOV of the focal layer (lines) and the localization precision measured at different positions of both illumination spots using the iSCAT signal from 40nm gold nanoparticles. The x-axis represents the distance from the center of the FOV, pointing out the homogeneity of the top hat illumination. The interference fringes show up since the line profiles were acquired on the “raw” iSCAT images before application of any median filter. For ease of interpretation, the line profiles are smoothed with a gaussian filter of 5 pixel radius. (b) Contrast of the iSCAT signal for the top hat illumination profile determined as (*I*_*background*_ − *I*_*particle*_)/*I*_*median filter*_ : frequency distribution (number or counts of respective contrast/grey values) for 683, 290 and 275 acquisition of 40nm (blue), 20 nm (red) gold nanoparticles (GNP) and PEG40K (green), respectively.

Besides widening the FoV, th homogeneous TH illumination resulted in an almost constant localization precision of individually scattering objects over the whole FoV. To determine the static localization precision, we prepared a sample of 40 nm gold nanoparticles (GNPs) coated with streptavidin, which we immobilized onto a coverslip via spin-coating, as established before [14]. We then observed the relative distance between two immobile particles over time. For this, we acquired a stack of 1000 images and extracted the positions of two GNPs using a localization algorithm described in the Methods section [40]. The standard deviation of the distance between the two objects was taken as an estimation of the localization precision of the iSCAT setup. These measurements lead to a constant sub-10 nm static localization precision throughout the FoV. In contrast, the Gaussian illumination profile resulted in a spatially strongly inhomogeneous localization precision that vastly deteriorated from around 5 nm in the focal center to above 15 nm around 30 µm away. For direct comparison, the localization precisions of both illumination profiles were acquired with a power density of about 2.3 *µW*/*µm*^2^ in the maximum.

A high frame rate is crucial for investigating sub-ms dynamics in biological sample, which is typically achieved by reducing the number of camera pixels and consequently of the FoV to a few *µm*^2^ only [12,41]. For our homogeneous TH profile and the instituted CMOS camera, we could achieve a frame rate in the order of 3000 fps with 300 *µs* exposure time. Of course, we could further enhance the frame rate by reducing the acquired region of interest. We have to note, that the limitation in the achievable sampling rates was only imposed by the characteristics of the detector at our disposal, and the setup may achieve higher performance with different devices.

We next tested the achievable scattering signal of our iSCAT setup, by monitoring and comparing the signal from immobilized 20 and 40 nm GNPs samples alongside 40-kDa PEG molecules, which represented more inert labels. For this, we determined the contrast levels (*I*_*background*_ − *I*_*particle*_)/*I*_*median filter*_ as the iSCAT signal *I*_*particle*_ over background signal *I*_*background*_ (with temporal median signal *I*_*median*_) for the TH profile for an acquisition time of 300 *µs* (Fig. 3.b), which highlight an excellent scattering signal even for the PEG molecules, scaling with the size of the molecules as expected. The contrast levels are influenced by characteristics of the detector such as full well capacity and may therefore be improved upon selecting further optimized detectors.

### 2.4 Single Molecule applications

We next demonstrated the performance and the sensitivity of our iSCAT microscope with TH illumination and the potential of correlative fluorescence recordings through single particle tracking (SPT) and TIRF measurements on model membranes. The supported lipid bilayers (SLBs) were generated from a 1:1 mixture of POPC (1-palmitoyl-2-oleoyl-glycero-3-phosphocholine) lipids and cholesterol through either spin-coating (Fig. 4a) or rupture of giant unilameller vesicles (GUVs, denoted GUV patches) (Fig. 4b) on Hellmanex-cleaned glass supports (Fig. 4a). They were stained with scattering DSPE (1,2-distearoyl-sn-glycero-3-phosphoethanolamine, 0.01 Mol%) and fluorescent DOPE (1,2-dioleoyl-sn-glycero-3-phosphoethanolamine, 0.01 Mol%) lipid analogues for iSCAT and TIRF recordings, respectively. For scattering biotinylated DSPE lipid analogues (DSPE-PEG2k-Biotin) were tagged with 40-nm large streptavidin-coated gold nanoparticles and for fluorescence DOPE tagged with the dye Atto488. With continuous iSCAT recordings (3000 fps) we could follow the spatio-temporal tracks of individual gold-nanoparticle tagged lipids with high spatial and temporal resolution over the whole FOV (Fig. 4a). In our measurements, the TIRF fluorescence signal guided the iSCAT acquisition towards the positions and dimensions of the GUV patches, as representatively highlighted in Fig. 4b for correlative iSCAT SPT and TIRF image data. As expected, the motion of the lipids was restricted to the GUV patches.

**Fig. 4.**
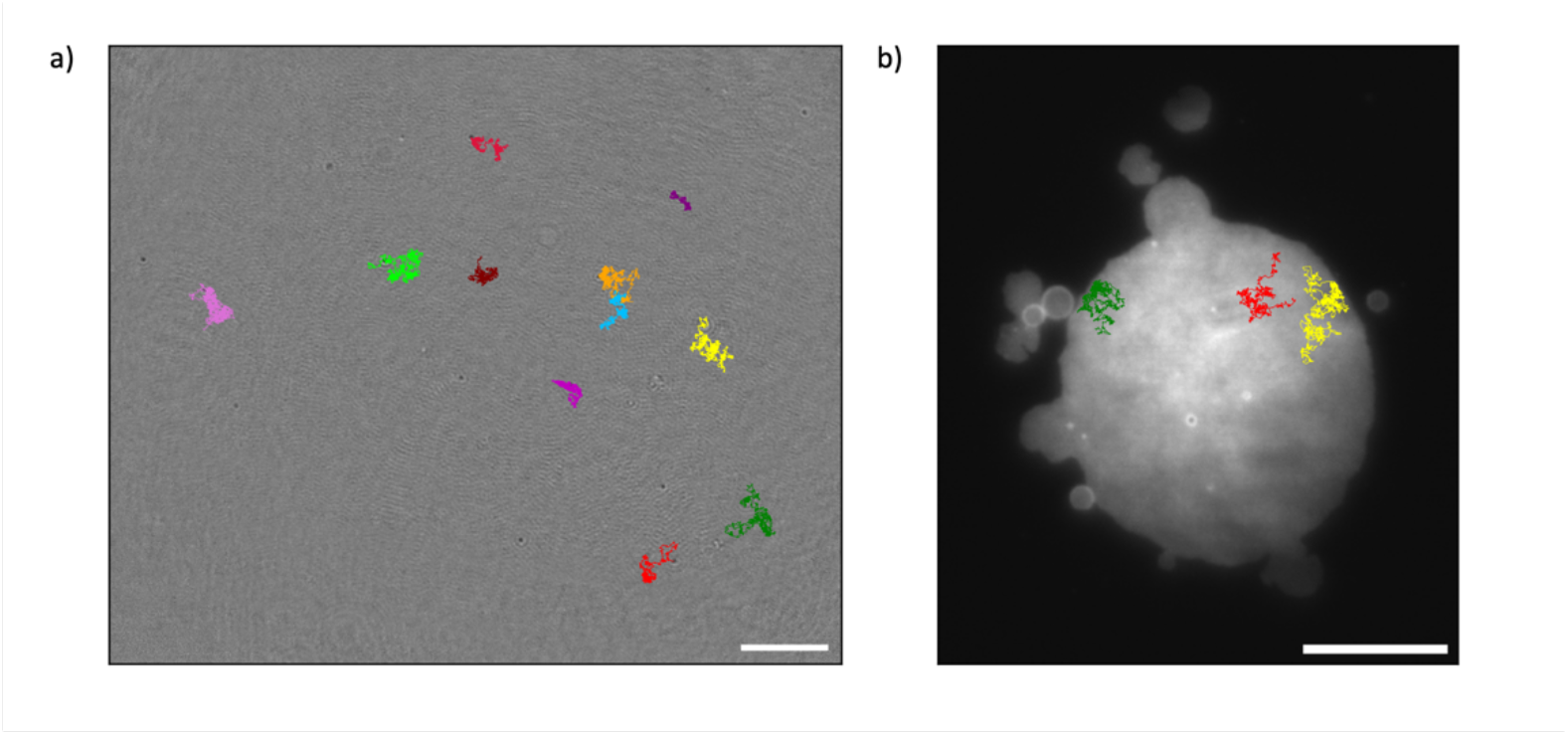
iSCAT-SPT data and correlative TIRF imaging. (a) Several representative tracks of individual 40 nm gold nanoparticles tagged DSPE-PEG2k-biotin lipid analogues on a POPC:Chol 1:1 SLB acquired at 3000 fps, and superimposed to an iSCAT image snapshot of the full iSCAT FoV (60×60 *µm*^3^). Scale bar 10 *µm*. (b) Representative TIRF fluorescence image of a GUV patch (labeled with fluorescent Atto 488 DOPE). Superimposed are representative tracks of 40 nm gold nanoparticles tagged DSPE-PEG2k-biotin lipids acquired with iSCAT, highlighting how the TIRF image is guiding the iSCAT recordings. Scale bar 5 *µm*.

To further outline the performance of our setup, we quantified the iSCAT-SPT data and thus the motion of the gold-bead tagged lipids in the SLB. For this, we followed a general procedure and calculated the mean-squared-displacement (MSD) of 128 individual tracks (Fig. 5a) and fitted them by a model of free diffusion, revealing the diffusion coefficient D alongside with a dynamic localization precision of *δ*. The curves did not show significant variation, and especially no strong deviation from a linear dependency, resulting in a distribution of values around an average of D = 0.30 ± 0.11 µm^2^/s (insert Fig. 5a) and δ = 20.1 ± 18.2 nm (Fig. S2). As a comparison, we also averaged all acquired 128 individual tracks to a single “ensemble” MSD curve, which was as well described by a model of free diffusion with the diffusion coefficient of *D* = 0.31 ± 0.0002 *µm*^2^/*s* and a dynamic localization precision *δ* = 18.7 ± 0.3 *nm* (Fig. 5a) This free, unrestricted diffusion of the target lipids is further highlighted in the inset of Fig. 5a, where we plotted the apparent diffusion coefficient D = MSD/4t against time t, obtained by dividing the average MSD curve by the time lag t and the dimensionality factor 4 (Fig. 5a, inset). The curve shows a constant diffusion coefficient over time, indicative of unbiased diffusion, while the dip at very short lag times stems from the motion blur typical of camera-based SPT measurements ([42,43]).

**Fig. 5.**
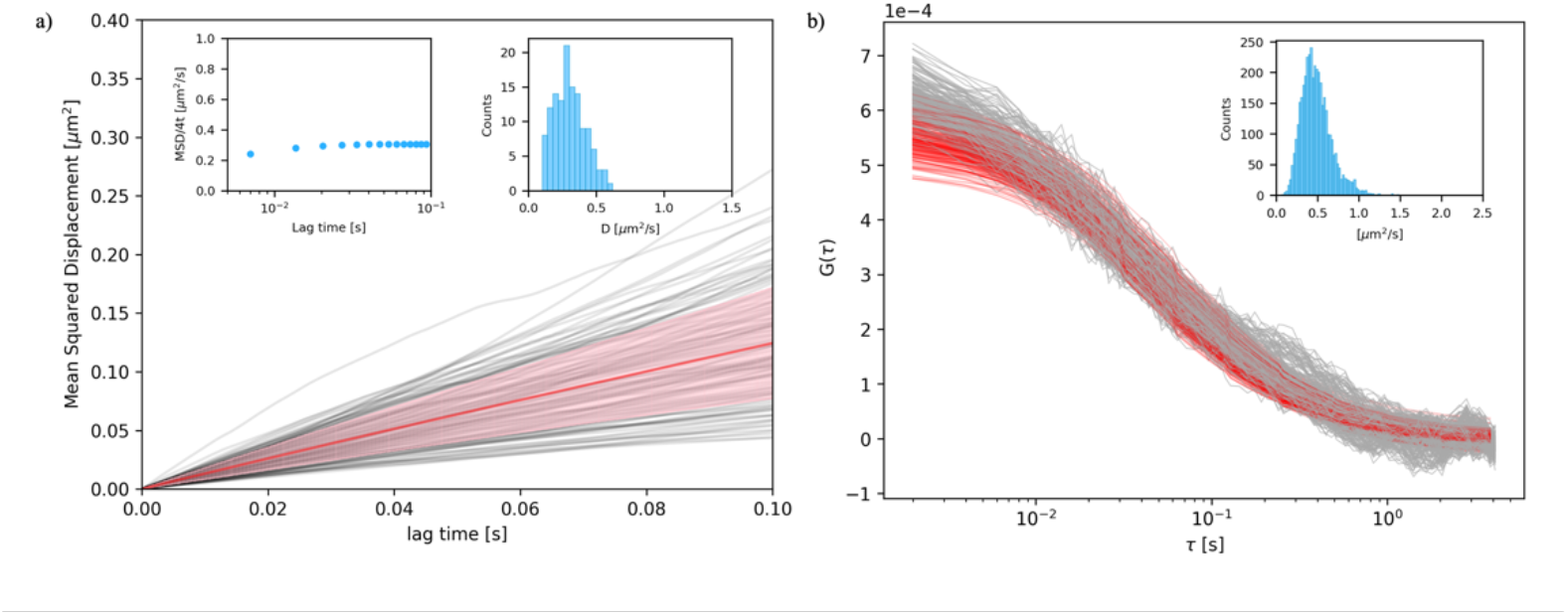
Complementary lipid mobility analysis with iSCAT-SPT and TIRF imaging FCS for the scattering and fluorescent lipid analogues diffusing in the SLBs (GUV patches) of Fig. 4 (POPC:Chol 1:1). (a) iSCAT-SPT data of the diffusion of the 40 nm gold nanoparticle tagged DSPE-PEG2k-biotin lipids: mean squared displacement (MSD) versus trajectory lag time t. The plot shows the MSD from 128 individual tracks (grey) together with the averaged MSD (ensemble with error bars from standard deviations of the mean in faint red). (Insert left) Plot of the ensemble MSD/4t, pointing out a free diffusion as a straight line. (Insert right) Frequency distribution of values of the diffusion coefficient D extracted from the MSD analysis of the 128 individual tracks. (b) Imaging FCS analysis: 455 ACFs curves (grey) together with their corresponding fits by Eq. S2 (red lines) of a representative TIRF image stack. (Insert) Frequency distribution of values of the diffusion coefficient D extracted for each individual ACFs fitting.

Through the TIRF channel of our setup we could further validate the homogeneous and free-diffusivity environment of the SLB patches. Recording the fluctuations in the fluorescence signal of the Atto488-tagged DOPE lipid analogues over time, we determined their mobility through imaging Fluorescence Correlation Spectroscopy (FCS). For this, we acquired nine different stacks of a 64×64 pixels region on the fluorescence-detection camera, depicting different 3.6 x 3.6 µm^2^ large areas on (different) SLB samples. Each stack was formed by 40,000 frames acquired with 2 ms exposure time. We subsequently binned each frame by averaging together the intensity collected from squares of 3×3 pixels within the 64×64 pixels, and performed the autocorrelation on the, in this way, jointed pixels of the stacks using the Fiji plugin for ImFCS [44]. For each stack we thus obtained a collection of 455 autocorrelation functions (ACFs, (64×64)/(3×3) ≈ 455). The 455 individual ACFs and their fittings are shown in Fig. 5b for a representative data set. We fitted a model of free diffusion (Eq. S2) [35,45] to each ACF of each data set, resulting in a total of 3967 values of the diffusion coefficient with a mean value and standard deviation of *D* = 0.49 ± 0.18 *µm*^2^/*s*. As expected, this value was slightly larger than that determined for our iSCAT-SPT analysis, since the gold-tagged lipid analogue displayed a slightly reduced mobility compared to the dye-tagged lipid analogue, as highlighted before [46–48]. Yet, most importantly both analysis approaches could be employed on the same sample and clearly depicted free diffusion.

## 3. Conclusions

In this work, we presented a new method to implement a custom optical setup with independent iSCAT and TIRF single molecule-sensitive channels. The setup we proposed here presents a number of advantages compared to similar protocols already present in literature [18]. First, we demonstrated the possibility of using true homogeneous illumination, that is, a top-hat shaped intensity profile of the incoming light on the sample, and still obtaining adequate imaging performance in iSCAT microscopy. We achieved such flat illumination profile using a refractive optical element, which was also presented in [27]. Yet, a similar flat illumination profile performance should also be realizable with other (sometimes more complex) approaches, as realized before for single-molecule based fluorescence microscopy (for example [49–52]). Second, we decoupled the TIRF illumination from the focus locking (as it was previously realized in [18]), thus making it easier to add more excitation wavelength for fluorescence and generally simplifying the device setup. Moreover, with a TIRF channel optimized for fluorescence imaging, rather than as a byproduct of the focus locking optical path, we showed how it is possible to attain single-molecule sensitive imaging FCS measurements, which may in turn be used to improve the quality of TIRF measurements [53] and complement the iSCAT data. Additionally, we provided a flexible and open-source software solution, based on ImSwitch, that can be adapted reasonably straightforward to other hardware configurations such as to support confocal imaging and related techniques [37]. ImSwitch is thus a viable solution for other implementations of iSCAT imaging Finally, we expect that the presented instrumental design will significantly lower the barrier for adoption of iSCAT microscopy by laboratories with the necessary resources and expertise, leading to a wider variety of application for this very promising microscopy technique.

## 4. Materials and Methods

### 4.1 Setup

We designed an ISCAT setup with a small footprint (1m x 1m), with the aim of being completely reproducible and straightforward to implement (Fig.1). All the optomechanical and optical components are commercially available and easy to source. For the iSCAT channel we used a single mode diode laser (iBEAM SMART 660, Toptica). The beam was directed through a half-wave plate (B. Halle) to linearly polarize the beam with respect to the optical axis [18]. To improve laser mode quality, we coupled the beam into a single mode polarization maintaining fiber (P5-630PM-FC-2, Thorlabs) and collimated the fiber output with a Plan Achromat Objective (4x, RMS4X, Thorlabs). A telescope system with magnification 1,25x was used to enlarge the beam diameter and meet the input requirements of the TopHat component (a|TopShape, Asphericon). The output of the TopHat component, that is, a collimated beam with uniform irradiance across its diameter, was focused on the back focal plane of a high-numerical-aperture objective (α Plan-Apochromat 63x/1.46 Oil Corr M27, ZEISS) with a 400 mm focal length (fl) lens (AC254-400-A-ML, Thorlabs). After the focusing lens, the beam was sent through a polarizing beam splitter (PBS, B. Halle) and a quarter-wave plate (B. Halle) and finally directed to the objective with a polarization-maintaining silver-coated mirror. The quarter wave plate circularly polarized the illumination beam. Due to the reflection at the glass-sample interface the circular polarization was inverted and, passing once more through the quarter wave plate, the light was again linearly polarized in the perpendicular direction to the illumination. Therefore, the PBS now reflected the signal towards the detection channel, where the signal was collected by a high-framerate CMOS camera (CB013MG-LX-X8G3, Ximea) to create an image. We focused the signal on the camera chip with a 400 mm-focal length lens (AC254-400-A-ML, Thorlabs) leading to an effective magnification of 153x, corresponding to a 90 nm effective pixel dimension.

For the TIRF channel we used a high-performance single-mode diode laser (Cobolt 06-MLD 488, Hübner Photonics). We coupled the laser in a single mode fiber (P5-488PM-FC-2) and collimated the fiber output with a Plan Achromat Objective (4x, RMS4X, Thorlabs). We focused the beam onto the back focal aperture of the objective with a 500mm-fl lens (AC254-500-A-ML, Thorlabs). We mounted the lens on a linear translation stage (XR50C/M, Thorlabs) enabling to switch between epi and total internal reflection illumination. The fluorescence signal was then collected by the objective and focused on the camera (Kinetix, back-illuminated Scientific CMOS (sCMOS), Photometrics) by a 300 mm-fl lens (AC254-300-A-ML, Thorlabs) leading to an effective magnification of 115x, corresponding to a 57 nm effective pixel dimension.

Appropriate filters were placed in front of the cameras in both the iSCAT and TIRF detection channels to prevent unwanted light leaks from the illumination channels.

Sample positioning and focusing was done exclusively through the stage, while the setup objective was mounted on a fixed optical cage system. Since the iSCAT signal relies on the difference in optical path between the scattered light from the sample and the back reflected reference beam, high stability was required hence the choice of the stage used to hold and move the sample became crucial. We employed a XYZ motorized stage (Mad-Deck, Mad City Labs Inc.) in combination with a Z-axis piezo nanopositioner for fine focusing (Nano-Z Series, Mad City Labs Inc.).

We integrated a focus-lock channel in the system to ensure a higher degree of stability in the axial direction of our sample, with a procedure similar to what reported in [37], which had the advantage of being readily available in the ImSwitch control software adopted for the control of the microscope [38]. To build this focus-lock channel we used a 780 laser (CPS780S) as probing signal. We cleaned the beam profile through a 50 *µm* pinhole located in the Fourier plane of a 4f system with 0.4x magnification. The collimated beam was channeled off-axis to the back aperture of the objective to achieve total internal reflection with a short pass dichroic mirror (BS HC750SP, AHF). The back reflection was picked up with a D-shaped mirror and sent to a CMOS camera (FLIR Blackfly S USB3.2 Cameras). The position of the intensity peak of the back reflection responded to a change in the axial direction of the sample. The position of the peak on the camera chip was recorded by a plugin in ImSwitch and, through a feedback loop, the z position of the sample was adjusted to keep the sample in focus using the Z-axis nanopositioning stage.

### 4.2 Microscope control software

The setup was fully controlled using ImSwitch [38], except for the Photometrics Kinetix camera in the fluorescence channel. We developed an ad-hoc version of it to allow a customized set of controls for the deployed cameras [54]. This version presented the following changes and additions: 1) a software interface to integrate the Ximea camera controls into ImSwitch; 2) camera acquisition running concurrently for each channel (iSCAT and focus-lock channel); 3) lasers controlled by the microscope package [55]; 4) stages controlled by the Python bindings of the µManager core [56]. The inclusion of these software packages provided a more flexible choice for reproducibility and deployment time optimization for the users. ImSwitch user interface (UI) provided the means to specify how to generate and apply the temporal median filter used on the iSCAT channel. The UI allowed to select: 1) the number of frames to record for the image stack that were employed for median calculation (by default this is set to 100 images); 2) the minimum stage step before collecting each image of said stack (by default this is set to 0.2 µm); 3) whether to apply the temporal median filter by division or subtraction of the incoming images. After selecting these options, ImSwitch was running a background thread to perform a loop which moved the X-Y µ-positioning stages in a square-like movement. For each direction, a number of steps equal to a quarter of the maximum number of frames was performed; for each step the routine saved an image from the iSCAT channel; at the end of this loop, the temporal median filter was calculated through NumPy. The ImSwitch live view command gave a preview of what was produced by applying in real-time the median filter to the collected images. After recording, if the temporal median filter was applied, the pixel data of the filter was stored together with the image stack in an HDF5 file.

### 4.3 Data analysis

To perform SPT on the iSCAT channel, after the acquisition of a stack of images, we applied the median filter dividing the raw images by the median filter previously generated in ImSwitch (Microscope control software). In order to filter out every entity that was not moving in the sample, we generated a median projection on Fiji software. Fiji generated a single image in which each pixel represented the median value of the single pixel over the whole stack. This image showed every object that was not diffusing in the sample. By means of an image subtraction we then ended up with a stack in which only the diffusing particles were highlighted. We then performed SPT and MSD analysis through a python code using trackpy package [40].

To perform imaging FCS, once we acquired a stack of 40000 frames in the TIRF channel, we uploaded the data in the Fiji plugin ImFCS [44]. This plugin performed the autocorrelation of each pixel’s signal throughout the stack resulting in as much ACFs as the number of pixel in the image. We applied a 5^th^ order polynomial photobleaching correction and we fitted each ACFs using the mentioned plugin [35] (Eq. S2).

### 4.4 Sample preparation

The supported lipid bilayer used in this study were prepared with two different techniques. To show the proof of principle of SPT in a large field of view, we spin-coated a mixture of POPC:Chol 1:1 (1-palmitoyl-2-oleoyl-glycero-3-phosphocholine, cholesterol, Avanti Polar Lipids) dissolved in 1:1 chloroform/methanol solution with the addition of 0.01 Mol % Atto 488 tagged DOPE (1,2-dioleoyl-sn-glycero-3-phosphoethanolamine, Atto-Tec) and 0.01 Mol% DSPE-PEG2k-Biotin (1,2-distearoyl-sn-glycero-3-phosphoethanolamine, Avanti Polar Lipids). 25 *µl* of the lipid solution (1 *mg*/*ml*) was dropped on a Hellmanex cleaned round cover glass (25mm diameter, 0.15mm thickness) and positioned on a spin coater. The cover glass was immediately spun for 30 s at 50 rotations per second such to enable the formation of SLB on the glass support. The cover glass was positioned in a liquid-tight chamber and 1 ml of phosphate buffer saline (PBS, 137 mM NaCl, 10 mM phosphate, 2.7 mM KCl) was pippetted on top of the glass. Afterwards, 5 *µl* of the 40 nm gold nanoparticles (GNPs) stock solution (BBI solutions, BA.SPT40/X, OD 10.1) was added to the sample and incubated for 10 minutes. Finally, we washed the sample to eliminate not bound GNPs.

To perform quantitative SPT measurements, the SLBs were formed through rapture of giant unilamellar vesicles (GUVs), denoted GUVs patches throughout. The same lipid solution described above was used to prepare GUV through electroporation. 5 *µl* of lipid solution (1 *mg*/*ml*) were dried on the wires of a GUV chamber, the GUV chamber was then filled with 400µl of sucrose solution (300 mM). The wires were connected to a function generator (PSG9080 Single Generator, Joy-it) to generate a sinusoidal alternating electric field with a peak-to-peak amplitude of 5.7 V at 10 Hz. After 1 hour the frequency was changed to 2 Hz for the following 30 minutes to allow the GUVs to detach from the wires. A round cover glass (25mm diameter, 0.15mm thickness) was treated through plasma cleaning to make the surface hydrophilic. The cover glass was then positioned in a liquid-tight chamber and 50 *µl* of GUV solution were mixed to 950 of PBS. Because the sucrose solution inside the GUV was denser then the PBS the GUV sank and due to the plasma cleaning, once they touched the cover glass, they raptured creating a homogenous SLB. The sample was washed to remove remaining floating GUVs. 1 *µl* of 40nm GNP (stock solution) was added to the chamber and let incubate for ten minutes. Finally, the sample is washed once again to remove unbound GNPs.

To prepare imaging FCS samples, we used a lipid solution POPC:Chol 1:1 (1 *mg*/*ml*). for GUV electroporation. After electroporation, we probed the SLB with Atto 488-tagged DOPE (50nM final concentration). We incubated for 10 minutes and then prepared the sample on a round cover glass as described for the SPT measurements.

## Acknowledgments

The authors greatly acknowledge financial support by the Deutsche Forschungsgemeinschaft (DFG, German Research Foundation; Germany’
ss Excellence Strategy – EXC 2051 – Project-ID 390713860; project number 316213987 – SFB 1278; GRK M-M-M: GRK 2723/1 – 2023 – ID 44711651; project PolaRas EG 325/2-1), the Free State of Thuringia (Advanced Flu-Spec / 2020 FGZ: FGI 0031; Multi-XUV / 2023 FGR 0054), the German Academic Exchange Service (DAAD, Graduate School Scholarship Programme, 2022, Project-ID : 57597951), the European Union’s horizon 2020 (research and innovation programme under grant agreement No. 779472), the Chan-Zuckenberg Initiative (napari Plugin Grants (Cycle 2), ID grant: DAF2022-311155), and PhotonHub Europe (Horizon 2020 research and innovation program, Grant Agreement n.101016665). Further, this work is supported by the BMBF, funding program LIVE2QMIC (FGZ: 13N15956) as well as Photonics Research Germany (FKZ: 13N15713 / 13N15717) and is integrated into the Leibniz Center for Photonics in Infection Research (LPI). The LPI initiated by Leibniz-IPHT, Leibniz-HKI, UKJ and FSU Jena is part of the BMBF national roadmap for research infrastructures.

## Data availability

Data underlying the results presented in this paper are not publicly available at this time but may be obtained from the authors upon reasonable request.

## Supplemental document

See Supplement 1 for supporting content.

