## Supplement Material for "Homogeneous large field-of-view and compact iSCAT-TIRF setup for dynamic single molecule measurements"

### 1. Interferometric stability analysis

The main, and only, objection found in relevant literature to the use of a large field of view for iSCAT microscopy is that the resulting interference originating from such an illumination scheme would cause the setup to not be "interferometrically stable" [1]. Our interpretation of this statement would be that the reflected field from the glass-sample interface would not be constant, hence rendering the background removal by median filter, as generally employed in iSCAT, to be impractical and degrading the final image quality. While the images provided with the manuscript should be sufficient, we opted to show that the setup presented is interferometrically stable with an explicit measurement. To this end, we prepared a sample containing clear PBS buffer and placed it on the microscope. We then acquired 10000 frames with an exposure time of 300 µs (for a total time of 3 seconds) of the full chip FoV. Using Fiji, we saved the line profile along the FoV in the perpendicular direction of the interference fringes for each frame of the stack (Fig. S1a). We subsequently calculated the autocorrelation of the intensity values of each of the 993 pixels of the line profile in time. We obtained that the average autocorrelation factor for each pixel was on average 0.99 (Fig. S1.b). The results enabled us to conclude that the setup was interferometrically stable, i.e. the interference fringes did not change in time, for far longer than the time required for each of the reported observations.

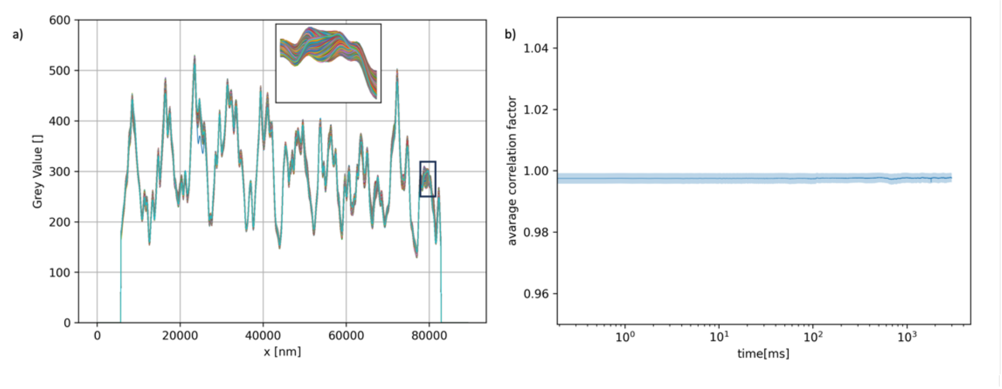


**Fig. S1** Interferometric stability of the iSCAT setup. (a) Superposition of 10000 line profiles (inset: zoom of the black frame in the main plot) of a raw iSCAT timelapse, traced perpendicular to the interference fringes elucidated in the main text. Each frame of the timelapse was taken with a 300us exposure time. (b) Average autocorrelation factor for the pixels in the line profile calculated over the entire timelapse in Fig. S1a. Each point in this line represents the average autocorrelation factor for each point in the raw iSCAT line profile. The shaded area represents the standard deviation of the average of every pixel at each time point.

### 2. Single particle tracking (SPT) analysis

To perform SPT analysis on the single particle tracks acquired we fit the Mean Squared Displacement (MSD) curves with the following model:

$$MSD(t_n) = 4t_n\left(1-\frac{2R}{n}\right)D + 4\delta_{xy}^2 \qquad \text{(Eq. S1)}$$

Where MSD($t_n$) is the mean squared displacement averaged in time, $t_n$ is the $n^{th}$ time interval, R is the motion blur correction factor, defined as the ratio between the exposure time and the frame time [2,3]), D is the artefact-corrected apparent diffusion coefficient, and $\delta_{xy}$ is the average dynamic localization precision. In our analysis we set R = (1/6)(300$\mu s$ / 303$\mu s$). We, then, restricted our analysis to truncated trajectory segments of 3000 localizations. The fitting is performed over the first 10% of localizations of each tracks [4].

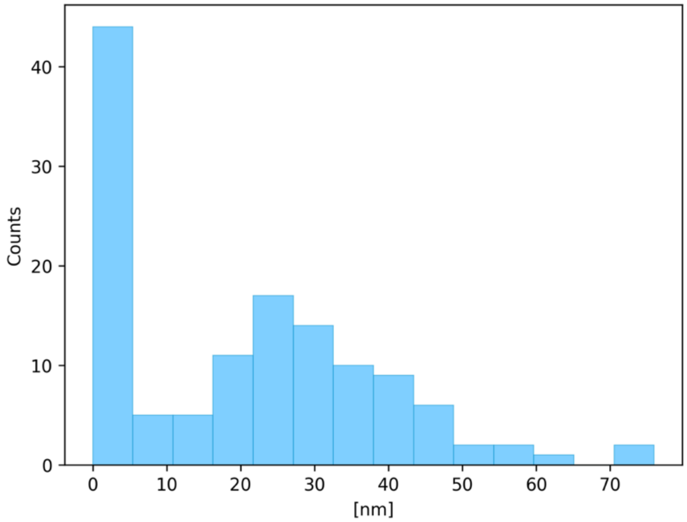


**Fig. S2** Frequency distribution of values of the dynamic localization precision extracted from the MSD analysis of the 128 individual tracks.

### 3. Imaging FCS analysis

To perform imaging total internal reflection FCS it is necessary to calculate the autocorrelation function (ACF) originating from the fluorescence intensity trace, and fit a specific model designed for imaging FCS [5,6]:

$$G(\tau) = \frac{1}{N}\left[\frac{\operatorname{erf}(p(\tau)) + \frac{(e^{-(p(\tau))^2}-1)}{\sqrt{\pi}p(\tau)}}{\operatorname{erf}(p(0)) + \frac{(e^{-(p(0))^2}-1)}{\sqrt{\pi}p(0)}}\right]^2 \quad where \quad p(\tau) = \frac{a}{\sqrt{4D\tau + w_0^2}} \qquad \text{(Eq. S2)}$$

Where D is the diffusion coefficient, $\tau$ is the lag time, $w_0$ is the radius $1/e^2$ of the lateral PSF (defined as $I = I_0 e^{-2(x^2+y^2)/w_0^2}$), $a$ is the pixel size of the used camera, and N is the average number of particles diffusing in the effective area. The observation volume, called as effective area in imaging FCS, is the result of the convolution between the pixel size and the PSF of the system. This quantity, in the case of ITIR-FCS, can be derived from the following formula, derived from [6]:

$$A_{eff} = \frac{a^2}{\left[\operatorname{erf}\left(\frac{a}{w_0}\right) + \frac{w_0}{a\sqrt{\pi}}\left(e^{-\left(\frac{a}{w_0}\right)^2} - 1\right)\right]^2} \qquad \text{(Eq. S3)}$$
